# Oxygen deprivation implicated in rapid coral mortality – an emerging perilous threat to coral reefs

**DOI:** 10.1101/2024.10.15.618297

**Authors:** Max S. Dhillon, Shannon G. Klein, Anieka J. Parry, Alessandro Moret, Carlos M. Duarte, Manuel Aranda

## Abstract

Tropical coral reefs are undergoing unprecedented degradation^1^, primarily due to the increasing intensity and frequency of marine heatwaves with climate change^2-4^. Coral bleaching is a well-known ramification of marine heatwaves, but rapid coral mortality is an emerging paradigm that visually manifests as the ‘sloughing’ of tissue from the coral skeleton. Unlike coral bleaching, coral tissue sloughing precludes any prospect of holobiont recovery beyond the initial onset^1,5,6^, indicating a life-or-death tipping point. Here, we experimentally confirm this phenomenon occurs when temperatures increase within temporal windows of hours to days, consistent with field observations^7,8^. Through microscale measurements of dissolved oxygen in the diffusive boundary layers of two abundant, keystone reef-building corals, we demonstrate that rapid temperature increases coincide with intrinsic oxygen deprivation, occurring before gross tissue disintegration or coral tissue sloughing. We propose that this distinct phenomenon arises from rapid heating, rendering the coral holobiont incapable of engaging in reactive processes to counteract the combined effects of heightened aerobic demands and impaired photosynthetic function. The passive diffusion of O_2_ from the surrounding bulk water is likely insufficient to meet the holobiont’s requirements, as explained by the Einstein–Smoluchowski kinetic theory of gases and Brownian motion^9,10^. These insights into coral tissue sloughing underscore the complexity of holobiont responses to stress and biophysical consequences of heatwaves. A granular understanding of these mechanisms is urgently needed, particularly regarding how heating rates may change under future climate scenarios, to re-evaluate the potentially under-recognised threats facing coral reefs.

## Main text

Over the past five decades, tropical coral reefs have endured unprecedented levels of degradation from pollution and climate change. Mass coral bleaching driven by intensifying marine heatwaves is the major driver of this degradation^2,3,11^, and involves the breakdown of the symbiotic relationship between the coral host and the endosymbiotic algal partner (family: Symbiodiniaceae). The breakdown of this relationship manifests in the obvious pale appearance of coral colonies and deprives the host of carbon and oxygen—the principal by-products of algal photosynthesis^12-14^. The scientific community has invested significant resources into understanding coral-bleaching mechanisms, their recovery processes, and the extent to which coral communities can adapt over time. However, there is emerging evidence of rapid coral mortality in response to recent, record-breaking marine heatwaves, without prior bleaching. These reports collectively describe a response where coral tissues visibly appear to slough or ‘melt’ off from the carbonate skeleton^6,8,15,16^. For example, dominant staghorn corals in the Caribbean have died abruptly from tissue loss, rather than from coral bleaching, following a heatwave across the Caribbean and eastern tropical Pacific^8^. While early reports of coral tissue sloughing originate from experimental studies in the early 1990s^17^, it was not until the 2015–2016 global bleaching event that observations of this phenomenon emerged on Australia’s Great Barrier Reef^3^. Further observations from eastern Australia^6^, southern Japan^7^ and the Caribbean^8^ are reported. With sea surface temperatures warming and marine heatwaves becoming increasingly intense^8^, mounting reports of this phenomenon may signify a concerning shift from mass coral bleaching to large-scale coral mortality, potentially indicating a ‘new normal’ for coral reefs in the face of climate change^1,8^.

With intensifying sea surface temperatures and increasing reports of coral tissue sloughing^6-8,15^, the presently poor understanding of the factors triggering it and the mechanisms underlying its manifestation pose a significant and urgent challenge. Interestingly, visual descriptions of coral tissue sloughing closely resemble the well-documented phenomenon of tissue disintegration in response to both environmental (extrinsic) and intra-tissue (intrinsic) oxygen deprivation. Tissue disintegration resulting from intrinsic hypoxia has been extensively reported across a wide range of organisms, spanning from complex species such as in humans^18-20^, rodents^19-21^, and fishes^19,22^, down to basal life-forms such as sea stars^23,24^. We therefore hypothesised that the rapid increases in temperature, and concomitant inhibition of the photosynthetic machinery of the algal symbiont^13^, lead to an intrinsic oxygen deficit—and physiologically underlies coral tissue sloughing (Figure 1).

**Figure 1:**
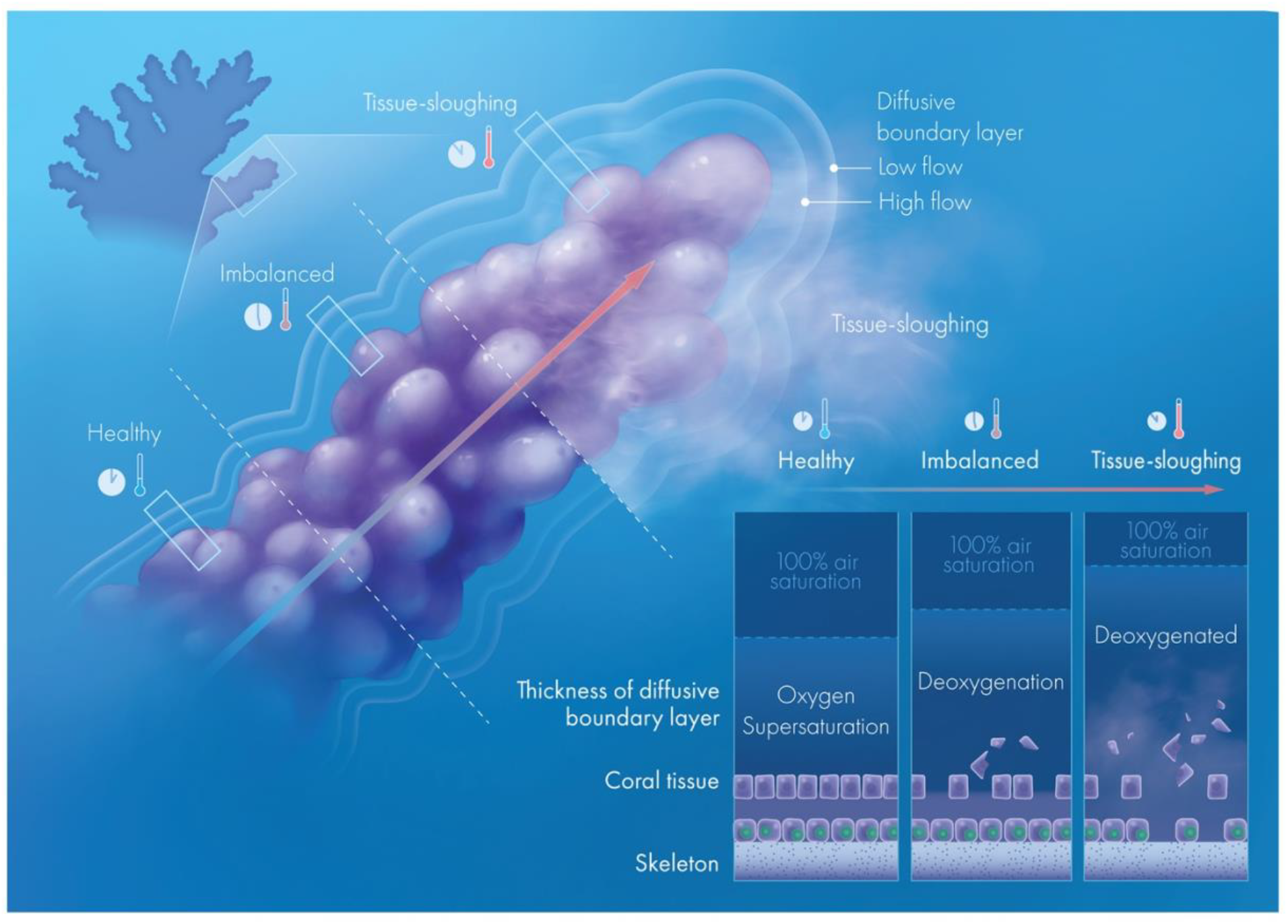
Conceptual representation of the coral tissue sloughing phenomenon. As the holobiont is exposed to thermally induced deoxygenation, tissue visibly starts to slough away from the skeleton. When conditions are optimal, a ‘healthy’ coral (Video 1) will be exposed to intrinsic oxygen supersaturation, as photosynthetic production of oxygen far exceeds the metabolic demand of the holobiont. At high temperatures, inhibition of the algal photosynthetic machinery leads to an oxygen deficit, as the sum of oxygen from the photosynthetic machinery and passive diffusion is not sufficient to maintain the aerobic metabolism of the holobiont. This leads to intrinsic deoxygenation (Video 2). If this imbalance persists, holobiont tissues will become intrinsically deoxygenated. At this point, coral tissue sloughing occurs, as the tissue disintegrates (Video 3).

Here, we present evidence from micro-sensor experiments showing that an internal oxygen deficit is indeed a precursor to coral tissue sloughing under rapid heating, from which corals cannot recover. Our findings confirm that short-term exposure to low oxygen levels reliably induces coral tissue sloughing, even in the absence of heat stress, further supporting the hypothesis that an internal oxygen deficit precedes this phenomenon. Importantly, these experimental responses resemble *in situ* reports of coral tissue sloughing and underscore the urgent need to re-evaluate the severity of climate change threats facing coral reefs (Figure 1).

### Acute heat stress induces coral tissue sloughing

Toward cataloguing and manipulating coral tissue sloughing at high resolution, we developed a bespoke, open, temperature-manipulation system (Supplementary Figure 1) to experimentally induce coral tissue sloughing in two reef-building corals by mimicking an acute heating event. *Acropora hemprichii* and *Stylophora pistillata* represent dominant members of reef communities across tropical reefs. To simulate quick and extreme changes to sea surface temperatures observed during especially severe marine heatwaves^25-28^ and associated *in situ* observations of coral tissue sloughing, we selected a heating rate of 1ºC per hour (Supplementary Table 1). This rate also mimics the thermal regime used by Baird et al. ^29^, where coral tissue sloughing was consistently observed. All coral fragments were exposed to the same thermal regime during a single photoperiod, starting at 2ºC below their summer mean and reaching 8ºC above (Supplementary Table 1). This regime induced coral tissue sloughing in 100% of the coral fragments from both species (*A. hemprichii n* = 96, *S. pistillata n* = 12). Consistent with *in situ* descriptions of coral tissue sloughing, all coral fragments showed signs of abrupt mortality, with tissue visibly ‘melting’ from the coral skeleton. Initial signs of coral tissue sloughing were subtle but rapidly progressed within minutes to hours (Videos 1–3), resulting in complete tissue loss within 24 h of the onset of coral tissue sloughing. These results confirm that rapid and intense heating are sufficient to induce coral tissue sloughing.

### Intrinsic oxygen deprivation precedes coral tissue sloughing

To test whether intrinsic deoxygenation precedes coral tissue sloughing, we integrated a microsensor set-up into our temperature-manipulation system to measure oxygen in the diffusive boundary layer of the corals. The diffusive boundary layer is a thin layer of seawater directly adjacent to the coral surface, wherein mass diffusion of oxygen (and other solutes) occurs between the coral surface and bulk seawater^30-35^. Changes in metabolic activity and intra-tissue oxygen levels are directly related to oxygen concentration gradients observed across the diffusive boundary layer^33,36-38^. Measurements of oxygen therein serve as a reliable proxy for changes in internal coral tissues, although changes in the diffusive boundary layer lag behind intra-tissue concentrations by seconds (see review by Nelson and Altieri ^39^). However, the extent to which the coral is isolated from the surrounding seawater (and the oxygen therein) is highly dynamic and changes according to numerous factors including water flow^40^ and the presence, and activity, of cilia^36-38,41^. We measured oxygen in the diffusive boundary layer by continuously logging oxygen at the coral surface and by measuring vertical profiles to examine changes in boundary-layer thickness.

At the surface of *A. hemprichii* fragments (*n* = 4), oxygen concentrations showed no discernible effect of individual genotypes across the 12-h experimental period (Figure 2D,E). At temperatures between 2ºC below (0 h) and 4ºC above (8 h) their summer mean, we observed concentrations consistent with hyperoxia (up to 750 μmol/L (approximately 400% air saturation); i.e., excess oxygen), indicating that supply of photosynthetic oxygen far exceeded demands of the holobiont (Figure 2F, region 1; no tissue sloughing observed). Hyperoxia within the diffusive boundary layer (and, by extension, within the coral tissues) persisted until a sharp and rapid decrease in oxygen occurred, which always preceded initial signs of coral tissue sloughing (Figure 2F, region 2; minor tissue sloughing observed).

**Figure 2:**
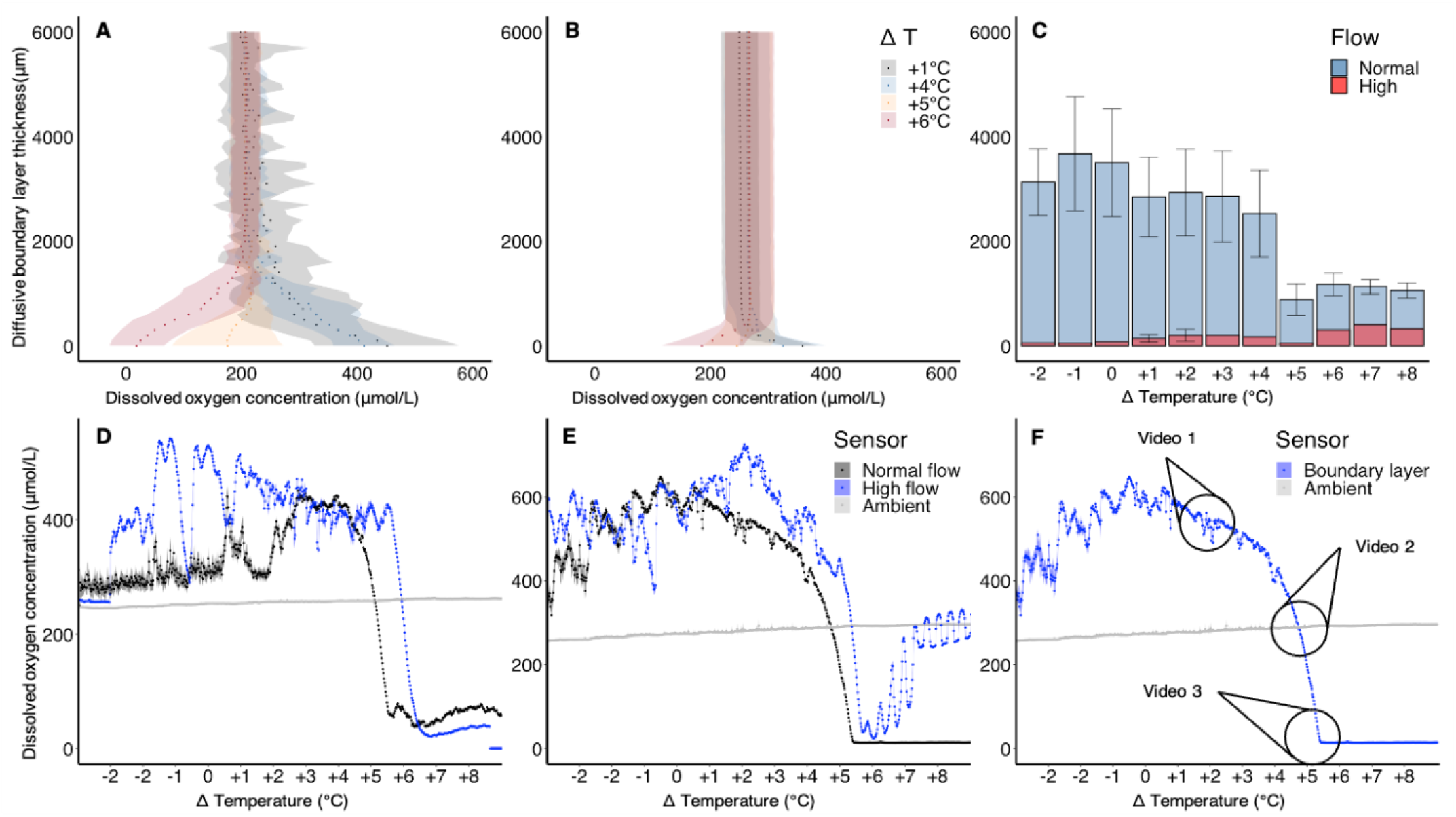
Spatial and temporal sequence of intrinsic oxygen deprivation that precedes coral tissue sloughing. **A**,**B**, Vertical profiles displaying oxygen concentrations from 10,000 μm from *Acropora hemprichii* surface to surface (x-axis; 0 μm) under normal-(**A**) and high-(**B**) flow conditions. Each point represents the average oxygen concentration across all profiles across all replicates for the corresponding temperature, with ribbons representing ±1 standard deviation. Delta temperatures represent the change in temperature from the summer mean. Temperatures plotted represent key points shared amongst all replicates: highest photosynthetic production to holobiont respiration ratio (i.e. highest holobiont excess oxygen), the beginning of imbalance, deoxygenation, and deoxygenated, respectively. **C**, Thickness of the diffusive boundary layer around *Acropora hemprichii* under normal-(blue) and high-flow (red) conditions. **D, E**, Oxygen concentrations in the diffusive boundary layer of *Acropora hemprichii* during an acute heat-stress event under normal flow (black) and high flow (blue) conditions for two genotypes (D and E). The oxygen concentration in the water column is depicted in grey (ambient). Each point represents the average dissolved oxygen concentration (μmol/L) across a minute of data, with ribbons representing ± 1 standard deviation. **F**, Three points correspond to the three timepoints of interest during an *Acropora hemprichii* exposed to an acute heat-stress event presented in Figure 1. At timepoint 1 (SVid 1), the coral is healthy and there is no sign of tissue sloughing. At timepoint 2 (SVid 2), subtle signs of tissue sloughing can be observed – the holobiont is reaching an imbalanced state, with oxygen concentrations in the diffusive boundary layer no longer exceeding that of the bulk water. At timepoint 3 (SVid 3), holobiont oxygen concentrations have collapsed, and severe tissue sloughing can be observed.

Oxygen concentrations eventually reached levels consistent with physiological definitions of severe hypoxia or anoxia (0–30% air saturation), at which point coral fragments died abruptly and showed severe tissue sloughing (Figures 2F, region 3; severe tissue sloughing observed). Vertical profiles of oxygen concentrations also showed stark shifts between oxygen production and consumption at temperatures 4–6ºC (8–10 h) above the summer mean (Figure 2A,B). The onset of these rapid oxygen declines across the diffusive boundary layers always preceded signs of tissue sloughing, linking reductions in intra-tissue oxygen levels with the initiation of coral tissue sloughing.

Measurements of algal operating photochemical efficiency (*F*_*q*_*′/F*_*m*_*′*) over the same time course indicated that the oxygen declines were largely caused by reductions in photochemical efficiency (Supplementary Figure 2). Given the extent and rapidity of the oxygen decline, the most plausible explanation is a collapse of photosynthetic oxygen production, likely compounded by substantially higher holobiont oxygen demands at elevated temperatures (Supplementary Figure 2). We hypothesise that this effect could be further exacerbated after the initial onset of tissue sloughing, as an influx of organic material into the diffusive boundary layer would fuel microbial respiration—a positive feedback loop known to occur in a heat-stressed sea star^23,24^.

To determine whether the observed data were specific to *A. hemprichii* or indicative of a broader phenomenon, we repeated these experiments with *S. pistillata*. Although the microenvironment of this *S. pistillata* exhibited slightly slower declines in oxygen levels, we observed the same pattern of decline into severe hypoxia/anoxia immediately prior to first signs of coral tissue sloughing (Figure 3). We hypothesise that morphological differences at the microscale might contribute to these differences. *S. pistillata* is less morphologically complex than *A. hemprichii* and has longer polyps that protrude farther from the coral’s surface, increasing surface area and access to O_2_ in the bulk seawater. Differences in metabolic activity between the two species may also contribute to explaining differences between species.

**Figure 3:**
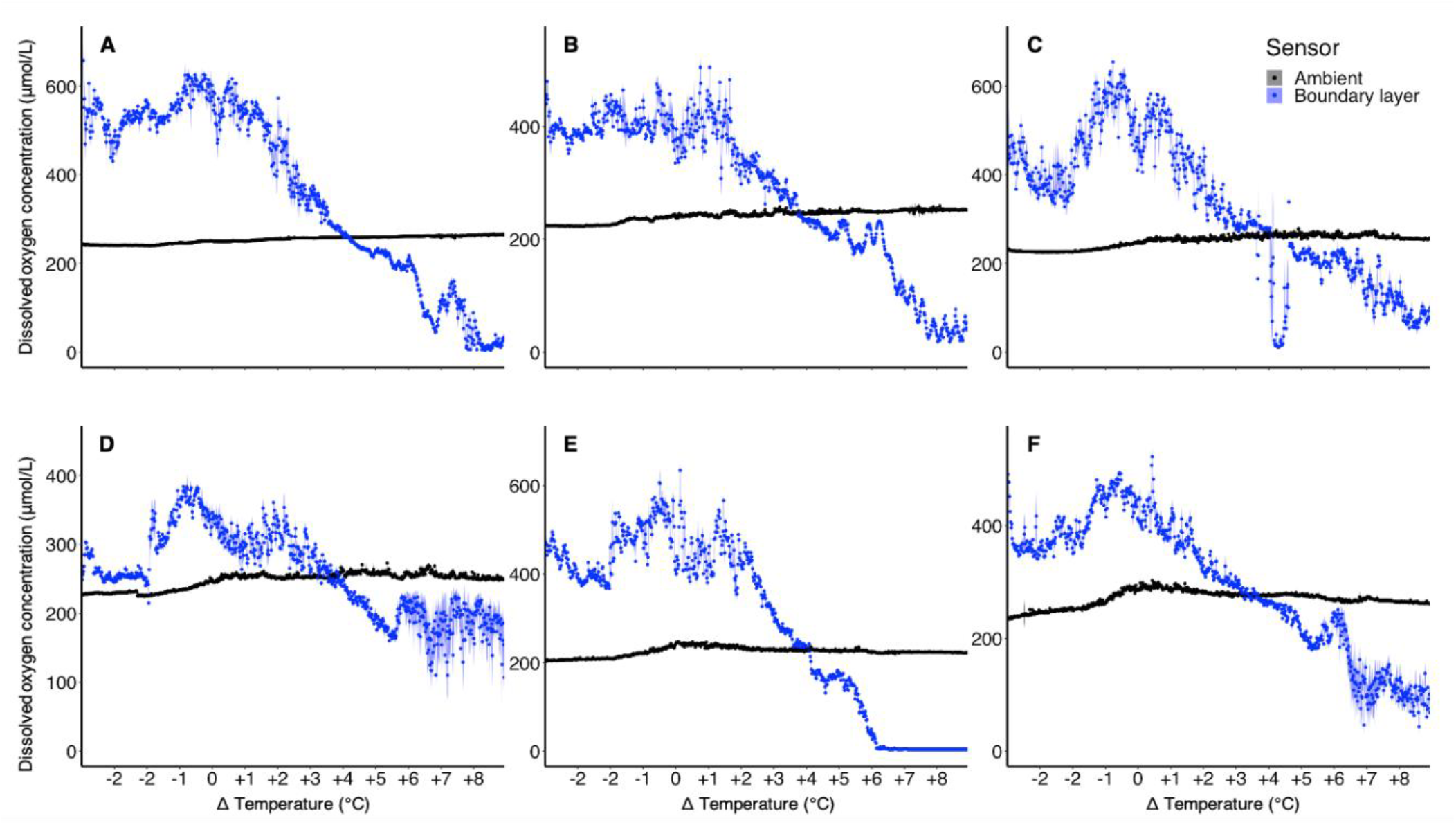
The coral tissue sloughing phenomenon is a multi-species response. **A-F**, Oxygen concentrations in the diffusive boundary layer of *Stylophora pistillata* through an acute heat stress event for six genotypes. The oxygen concentration in the water column is depicted by black (ambient). Each point represents the average dissolved oxygen concentration (μmol/L) across a minute of data, with ribbons representing ±1 standard deviation. Delta temperatures represent the change in temperature from the summer mean.

The role of higher flow rates in mitigating the effects of heat stress on coral bleaching is well-documented^42-44^. An increase in flow is anticipated to result in a thinning of the diffusive boundary layer and increase solute mixing. Both factors contribute to an enhanced availability of oxygen from the water column. The thinning of the diffusive boundary layer reduces the distance that molecular oxygen must passively diffuse before reaching the coral surface (Figure 2C). This is crucial because, according to the Einstein–Smoluchowski diffusion equations, the root mean square displacement (*λ*) of diffusing molecules is proportional to the square root of the product of time (*t*) and the diffusion coefficient (*D*) (*λ* = √2*Dt*). For example, at a temperature of 30ºC and a salinity of 40 psu, a molecule of oxygen will diffuse 0.0711 mm in 1 s, 0.711 mm in 100 s, and 7.11 mm in 10000 s. A small decrease in diffusion distance will thus result in substantially less time required for oxygen to reach the coral surface (due to the square root dependence on time), thereby enhancing oxygen availability and supporting critical metabolic processes in the coral tissue. We thus sought to ascertain if higher flow rates postpone the onset of coral tissue sloughing. Our results show that elevated flow rates delayed the onset of internal oxygen deprivation by over 1°C in *A. hemprichii* (Figure 2D,E).

These data link oxygen deprivation to the coral tissue sloughing response. We next tested whether oxygen deprivation could induce coral tissue sloughing in the absence of thermal stress by subjecting both species to closed-chamber incubations in complete darkness under ambient temperature. We hypothesised that, by removing the supply of oxygen from both photosynthetic production and passive diffusion from the surrounding seawater medium, we would observe coral tissue sloughing once the water had reached anoxia. All fragments of both species (*A. hemprichii* n = 9, *S. pistillata* n = 9) displayed the same coral tissue sloughing response after 4 h of anoxia, providing additional support for a role of intrinsic oxygen deprivation in triggering coral tissue sloughing (Supplementary Table 2).

We found no experimental reports of tissue disintegration in response to intrinsic oxygen deprivation in corals, however, one study reported instances of tissue sloughing in aquarium-reared corals when oxygen values in the diffusive boundary layers were near zero under heat stress^16^. While no other studies measured oxygen in the diffusive boundary layers of corals under warming, *in situ* observations of an acute deoxygenation event in Panama have linked coral tissue loss and mass coral mortality to extrinsic oxygen deprivation^15,45^. Our results also align with observations of tissue loss in the centre of coral colonies after unusually hot and low-flow conditions in waters off southern Japan^7^. The authors hypothesised this response was indicative of anoxia. In an examination of 11 instances of hypoxia-induced mass mortality within atoll lagoons in the Pacific, 10 of these events occurred either during, or at the conclusion of, the warm season—implicating a role of greater oxygen demand under higher sea surface temperatures^46^. *A. millepora* experiences complete tissue loss and subsequent mortality upon acute exposure to elevated temperatures and inhibition of photosynthesis by treatment with the herbicide diuron. This occurred despite the presence of ample oxygen availability (100% air saturation) in the surrounding seawater^47^. Under thermal stress, corals in high-flow environments survive better than those in low-flow environments^42^. Taken together, and in line with our findings, these results suggest that high-flow conditions and access to oxygen could alleviate and/or delay tissue loss by reducing diffusive-boundary-layer thickness in a manner like how high flow alleviates coral bleaching under heat stress.

### Coral tissue sloughing is the nexus of acute heat-stress and oxygen deprivation

Corals naturally experience low intra-tissue oxygen levels in darkness when Symbiodiniaceae spp. cease photosynthesis^32,48^. However, this can also occur in the daytime when conditions are not conducive to photosynthesis (such as during thermal stress), as the coral holobiont shifts from a net producer of oxygen to a net consumer^49,50^. The symbiotic relationship between the coral host and other holobiont compartments instead tips towards competition for the limited oxygen supply that can be extracted from the bulk water^39^. We observed that photobionts survived and were still capable of photosynthetic oxygen production after the corals had undergone tissue sloughing in response the acute thermal regime (cf. Methods). In cases where photosynthesis is inhibited during the daytime—as is typical during marine heatwaves—light-enhanced respiration likely exacerbates holobiont oxygen demand, and likely also competition for oxygen among holobiont compartments, as holobiont respiration rates can be between 6- and 12-fold higher than at night^33,51,52^. Strong evidence demonstrates that coral holobionts can regulate respiratory rates to conform to reduced oxygen availability^32,33,35,39,53,54^. Yet, elevated temperatures escalate the respiratory demands of the coral holobiont^50,55-57^, while also reducing oxygen availability through reductions in oxygen solubility^58,59^.

Like complex metazoans, shallow-water corals detect and react to low-oxygen conditions via gene networks involved in hypoxia-stress mitigation^60,61^. A recent study investigated the interaction between heat stress and extrinsic deoxygenation on corals and found similar stress responses to heat and extrinsic deoxygenation^62^. Akin to extrinsic deoxygenation, acute heat stress induces the transcription-factor gene *Hypoxia Inducible Factor* (*HIF-1*), indicating that corals respond to acute heat stress by initiating a hypoxic response. When heat stress and extrinsic deoxygenation co-occur, the cumulative oxidative stress from both factors may hinder HIF activity and the hypoxia-response system^62^. Consequently, the observed combination of rapid heating and intrinsic deoxygenation might hinder the corals’ ability to mitigate acute stress through reductions in aerobic respiration and the prevention of metabolic collapse. While the temporal scale of the HIF and hypoxia-response system in corals is not well understood, another plausible explanation for coral tissue sloughing could be their limited capacity to sense and respond to intrinsic oxygen deprivation over short time frames. Investigations into the molecular responses of corals during acute thermal regimes that induce coral tissue sloughing thus warrant immediate attention.

### Implications of coral tissue sloughing for coral-reef research

Coral tissue sloughing as we describe is clearly distinct from coral bleaching, but its visual appearance at a macro-scale could easily resemble the aftermath of coral bleaching or coral disease. Current limitations in the temporal resolution of reef monitoring across the majority of the world’s reefs often lead to coral bleaching or coral mortality as the only reliable metrics. Consequently, we believe that misdiagnoses may exist in previous data collections, wherein mortality has been attributed to a lack of recovery after coral bleaching when instead it may have derived from coral tissue sloughing. In line with observations from Hoegh-Guldberg *et al*. ^8^ of staghorn corals dying “abruptly”, Leggat *et al*. ^6^ describe “severe-heatwave-induced mortality events” characterised by rapid reef decay at a rate far higher than that observed after typical bleaching events, which usually show no mortality until 4–6 weeks following the onset of the event^5^. Hughes *et al*. ^1^ also reported “whole-colony mortality” in which corals appeared “white with fully intact fine-scale skeletal features”; a similar condition to what we observe following coral tissue sloughing. We thus posit that coral tissue sloughing likely underlies at least some records of coral mortality and emphasises the need for targeted efforts to observe distinct coral responses during intense marine heatwaves. Such efforts would help to understand if shifts from mass coral bleaching to large-scale coral mortality could become a new reality as marine heatwaves become more extreme^1,6-8^.

Although our study did not investigate the thresholds of heating rates that trigger coral tissue sloughing as opposed to coral bleaching, the short thermal regime examined is comparable with acute heating during modern-day marine heatwaves. Current stress-monitoring metrics, like degree-heating weeks, are designed to predict mass coral bleaching that occur in response to chronic temperature stress. Such metrics combine both the intensity and duration of heat stress events in singular metrics, challenging their ability to detect peak temperatures and rates of heating^63,64^—the crucial variables in predicting this response. There is a clear need for improvements in the spatial and temporal resolution of stress metrics and sea surface temperature monitoring to better capture coral-reef weather, potentially reaching sub-daily timescales to identify conditions that induce coral tissue sloughing. Simultaneous efforts to distinguish between coral bleaching and coral tissue sloughing, both in field and experimental studies, will be a crucial first step. Suggestions for enhancements in both spatial and temporal resolutions of reef monitoring, exploiting advancements in robotics and artificial intelligence, represent a promising avenue.

If large-scale mortality events on coral reefs prevail, all aspects of coral-reef research must adapt. Updating models that project future coral-reef responses to recognise and predict mass-mortality events, rather than just bleaching, will require considerable amounts of data. This will not only require greater temporal and spatial resolutions of sea surface temperatures, but also an understanding of which stress metrics and/or marine-heatwave characteristics reliably predict this response. The next step could involve identifying different traits that could help safeguard corals against tissue sloughing. For example, research streams presently focusing on mechanisms to mitigate the impacts of climate change on coral reefs, including assisted evolution and coral probiotics^65^, might also have to split their focus between increasing coral resilience to bleaching and tissue sloughing. Recognising the enormity of this challenge, collaboration amongst physiologists, ecologists and climate and ocean scientists will help to understand what data are required to meet this challenge.

## Acknowledgements

We are grateful for the support offered by the Coastal and Marine Resources (CMR) Core Lab at the King Abdullah University of Science and Technology; specifically, to H. Abdulbaqi, Z. Batang, N. M. Alikunhi, and F. B. Lampos. Figure 1 was created by X. Pita, Creative Lead for Research, with creative input from Heno Hwang, both from the Research Communication department at the King Abdullah University of Science and Technology.

## Author contributions statement

M.A conceptualised the idea. M.S.D, S.G.K, and M.A conceptualised and designed the research. M.S.D designed and fabricated experimental system. M.S.D and S.G.K developed methodology, with contributions from A.J.P. M.S.D and S.G.K conducted the experiments, with contributions from A.J.P and A.M. M.S.D performed data analyses, with support from S.G.K and A.J.P. M.S.D, S.G.K, and M.A interpreted the results. M.S.D wrote the initial draft, with major contributions from S.G.K. M.S.D, S.G.K, and M.A reviewed the draft, with contributions from C.M.D. M.A and C.M.D acquired funding and resources. All authors edited and approved the final manuscript.

## Data availability

Data generated by this study, supporting Figures 2-3 and Supplementary Figure 2 are available in Source Data files 1-8. All data generated by this study will be deposited in Dryad.

## Code availability

The R script needed to produce the analysis will be deposited in Dryad.

## Supplementary materials

Materials and methods Supplementary figures 1 and 2

Supplementary tables 1 and 2

Movies 1, 2 and 3

