## Supplementary information (methods, supplemetary figures 1 and 2, supplementary tables 1 and 2) for "Oxygen deprivation implicated in rapid coral mortality – an emerging perilous threat to coral reefs"

### Materials and Methods

#### *Study species and collection*

*Acropora hemprichii* and *Stylophora pistillata* were selected due to their abundance on global reefs, importance as dominant reef builders and their role as focal species in published empirical studies[1]. Approximately one third of each colony was sampled, to limit damage to the donor colony. Colonies were sampled at least 5 m apart to maximise the likelihood that the colonies were of different genotypes[2]. Collections were completed in June 2021, after which each colony was labelled, wrapped in bubble wrap, and placed in a cool box of seawater. The seawater in the cool box was constantly monitored and water changes were made intermittently during transportation to the laboratory facilities. The corals were placed into 150 L indoor aquaria supplied with a continuous flow of filtered seawater (flow rate ≈ 120 L/h) under a 12 h light:12 h darkness photoperiod using two Phillips CoralCare Gen2 LED lights (~350 μmol photons $m^{-2}$ $s^{-1}$ at the surface of the coral). The setup maintained the corals at conditions (temperature, salinity and pH) resembling field conditions of the donor reef (28ºC, 40‰ salinity, ~350 μmol photons $m^{-2}$ $s^{-1}$). To obtain fragments of a suitable size for experimentation, we used Aquamedic Coral Cutters (moon type) to separate small fragments (~5 cm branches) from the donor colony. All fragments were allowed a minimum of 4 d recovery prior to experimentation.

#### *Experimental system*

A bespoke aquaria system was constructed for this study, which comprised two small chambers (~1 L) held within a larger runway aquarium that acted as a water bath (~100 L; Supplementary Figure 1). The two chambers were fed with fresh seawater via a large sump (1,000 L) at a rate of 250–300 mL/min, equating to complete water exchanges once every 3 min. The outflow of each 1-L aquarium was discarded to ensure independence. We used six 650-W titanium heaters (D-D The Aquarium Solution Ltd) controlled by dual digital-temperature controllers (D-D The Aquarium Solution Ltd) in the 1,000 L sump and used four 200-W titanium heaters (D-D The Aquarium Solution Ltd) and the same temperature controllers in the runway aquarium to minimise heat loss. A large, commercial air pump (Linear Air Pump ET 100 80 W, Charles Austen Pumps Ltd) maintained dissolved oxygen concentrations at ~100% air saturation in the 1,000 L sump.

In each experiment run, we used two chambers; one that contained a coral fragment allocated to microsensor measurements (chamber 1), and the other that contained a coral of the same genotype, which was allocated for measurements of photochemical efficiency (chamber 2, see Symbiodiniaceae photochemical-efficiency section). Prior to the commencement of each run, the coral fragments were given 4 h to acclimatise to the chamber conditions at an ambient temperature of 2ºC below the summer mean. The temperature was held constant at 2ºC below the summer mean for 1 additional hour and then increased by 1ºC per hour until the nominal temperature had reach 8ºC above the summer mean (see Supplementary Table 1). In another experimental study, this rate of temperature increase had been shown to result in tissue sloughing[3].

*Microsensor measurements and approach*

We used two Unisense Clark-type OX-50 microsensors (tip size: 40–60 µm) simultaneously for oxygen measurements; one in the diffusive boundary layer and one in the water column to measure ambient oxygen. Prior to each experiment, a two-point calibration was performed by using 100% air-saturated fresh sea water (achieved using an air bubbler for 10 min) of known salinity and temperature and a solution of sodium sulfite ($Na_2SO_3$) in fresh seawater (0% oxygen) [4,5]. The microsensors were positioned using a MicroProfiling System (Unisense A/S), which includes two motor controllers to allow precise (0.5-µm increments) movements in the X and Z axis. A UniAmp Multi Channel (Unisense A/S) connected to and recorded data from the microsensors and SensorTrace Suite (current version: v3.4.400; Unisense A/S).

*Microsensor-placement protocols*

All microsensor measurements were taken adjacent to the surface of the coenosarc of an upwards facing, mature part of the coral fragment. This meant that the microsensors were not placed near areas of the coral that may naturally have a lower symbiont density, such as newer tissue close to the tip, or parts of the colony facing away from the sun/aquarium lights. Two methodologies were used to collect data with microsensors: one being through holding microsensors at the surface of the coral to collect data on oxygen dynamics within the DBL; and the other being through 10-mm profiles to observe diffusive boundary-layer dynamics at different temperatures. In all instances, if biofouling of the microsensor (through mucus or algae interference) was suspected, if a microsensor was broken or incorrectly calibrated, or if the software stopped working properly, the experiment was halted and reset. The two microsensors were placed concurrently, with one in the bulk water and the other at the coral surface. For experiments focused on oxygen dynamics within the diffusive boundary layer, sensors were held at the corals surface for the entirety of the acute heating simulation. For the diffusive boundary-layer profiles, the depth of the corals' surface was recorded and used as the endpoint. The start-point was set at 10 mm away from the surface, and the micromanipulator was programmed to automatically move toward the coral in 100 μm steps. At each step, the microsensor was programmed to wait for 1s, and then measure for 10s.

*Symbiodiniaceae photochemical efficiency*

Operating photochemical efficiency ($F_q'/F_m'$) values were determined using pulse-amplitude modulation fluorometry (PAM) using a MINI-PAM-II/R (Walz) equipped with 5.5-mm fibreoptics. $F_q'/F_m'$ measurements were conducted hourly on light-adapted coral fragments precisely before each incremental temperature increase during the heat-stress experiments. Acknowledging the potential interference of the fibre-optic cable with the corals' diffusive boundary layer, $F_q'/F_m'$ assessments were carried out on separate coral fragments of the same genotype. These $F_q'/F_m'$ coral fragments were housed in chamber 2 positioned adjacent to the chamber containing the coral fragments in chamber 1, wherein microsensor measurements were performed. Both sets of corals (in chambers 1 and 2) were subjected to the same heating regime over identical timeframes.

*Flow experiments*

The high-flow experiments adhered to the same thermal regime outlined above (Supplementary Table 1) and employed the same two methods for measuring oxygen in the diffusive boundary layer. Conditions within the coral-fragment chambers mirrored those in the ambient-flow experiments, with the addition of a single aquarium circulator pump in each chamber (6-W AC submersible pump). These pumps induced seawater turbulence exceeding 10 cm/s. The water-current velocity was carefully selected to significantly diminish the size of the diffusive boundary layers without displacing the fragments or affecting microsensor measurements. Seawater replenishment rates (250–300 mL/min) remained consistent between the high and low flow experiments.

*Extrinsic oxygen-deprivation experiments*

Closed-chamber respirometry experiments were performed in a setup comprising 12 sealable glass chambers (290 mL volume, Weck®, Germany), following ref. [6]. Each chamber was fitted with a fibre optic oxygen-sensor spot (PyroScience), in accordance with the manufacturers' recommendations, and a fibre optic cable (PyroScience) was positioned over the sensor spot from the outside of the chamber wall. Each cable was connected to, and communicated with, a Firesting Pro® 4 oxygen logger (PyroScience) via a PC, which equated fluorescence measurements from the sensor spots to concentrations of dissolved oxygen using commercial software and compensated for temperature and pressure (PyroScience). The sensor spots were calibrated using a two-point calibration (100% air saturation, and 0% oxygen via $N_2$ gas displacement). Each chamber contained a pre-sterilized, 3D-printed base that was specifically designed so the coral fragments were suspended via magnets through the wall of the chamber, which was capable of rotation and movement. The chambers were filled with autoclaved seawater to minimize background metabolism of the bulk seawater. All chambers contained a 3-cm stirring bar and were placed on a magnetic stir plate (IKA™, RT 15) that was set to 300 rpm and 2°C below the summer mean. The setup was placed inside a Percival incubator (Percival Scientific, model: I-22LLVLC8) that controlled atmospheric temperatures at 2°C below the summer mean and maintained complete darkness.

Two experiments were conducted, one for each species, and consisted of three coral fragments per genotype (n = 9 fragments per species) and three seawater blanks (n = 3). The time at which each chamber reached anoxia (0 mg $O_2$/L) was recorded, and each coral fragment was removed after exactly 4 h of exposure to anoxia. The corals were carefully inspected and changes to the coral tissue and water quality were compared to observations of the same fragment prior to the commencement of the experiment. Oxygen concentrations were recorded in the seawater blanks until all coral fragments were removed from the experiment. After each coral fragment was removed from their respective chamber, they were photographed and immediately transferred to holding aquaria for recovery. The recovery aquaria maintained the temperature at 2ºC below the summer mean and oxygen at 100% air saturation. Each coral fragment underwent a 24-h recovery period in the aquaria (Supplementary Table 2).

*Symbiodiniaceae cell survival*

At the conclusion of the heat-stress experiments, coral fragments were moved into 10 mL of fresh autoclaved seawater, and the remaining tissues were airbrushed from the skeletons. Tissue lysates were homogenised for 7 sec (T-18 Ultra Turrax, IKA, Germany) and then centrifuged for 5 min at 500 *g* (Centrifuge 5430 R, Eppendorf, Germany) to separate the symbionts from the host components. The symbiont pellet was then resuspended in 10 mL of autoclaved microfiltered seawater and incubated in small chambers connected to the same PyroScience oxygen-monitoring system. Samples were moved into temperature- and light-controlled incubators (Intellus Control System, Percival, Iowa, US), with temperature and light set to be the same as the holding tanks. Oxygen concentrations were continuously measured firstly for 30 min in light, and secondly for 30 min in the dark. This revealed a source of photosynthetic oxygen production, as rates of oxygen consumption were lower in the presence of PAR. These results confirm that at least some symbiont cells were alive in the sampled tissue following mortality of the coral host.

*Statistical analyses*

To ascertain the median effect dose ($ED_{50}$) corresponding to a 50% decline in operating photochemical efficiency ($F_q'/F_m'$), we employed three-parameter dose-response models (Supplementary Figure 2). The analyses was conducted using the ‘drc’ package in R (v4.2.1) [7]. $ED_{50}$ is frequently employed as a metric in coral thermal-stress assays, serving as a sub-lethal thermal threshold[8-10].

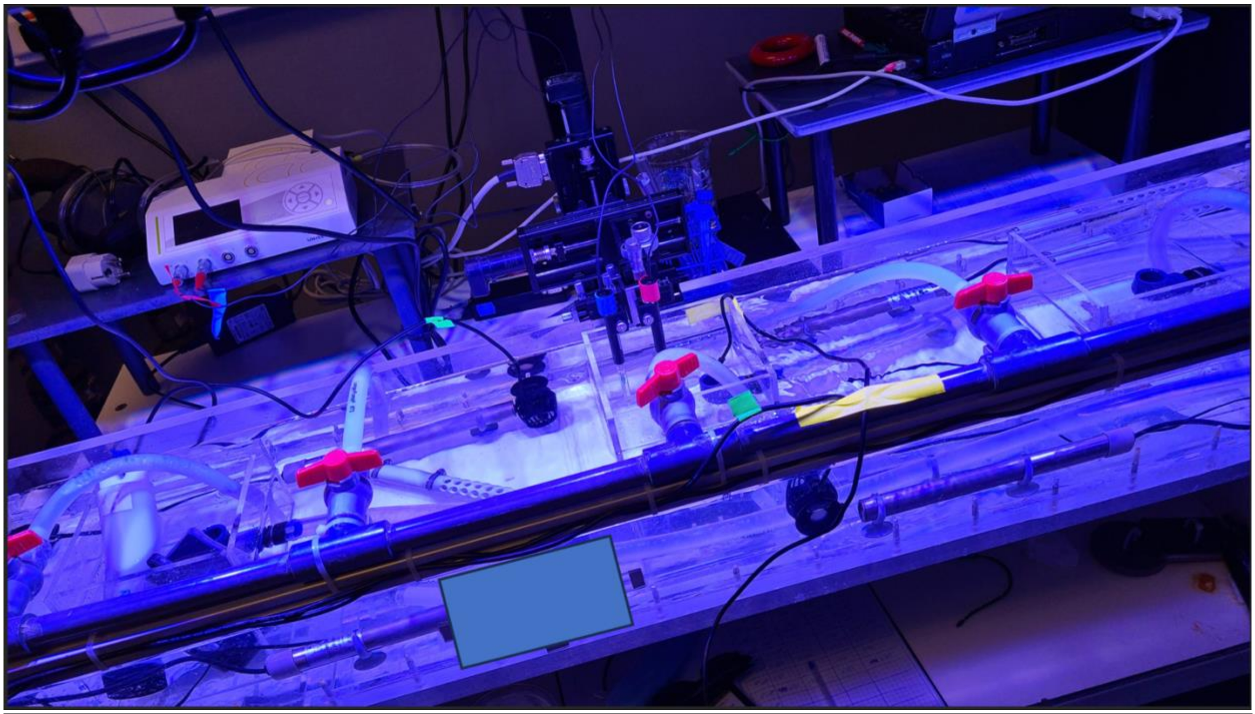

**Supplementary Figure 1: Bespoke temperature manipulation system.** A top view of the experimental setup, including the water bath, the two chambers, and the microsensor micromanipulator arm behind. Each chamber was subjected to the same flow of temperature-manipulated seawater.

**Supplementary Table 1: Temperature regimes across the entire study.** The bespoke experimental system was designed to reduce temperature variance at each setpoint among replicates, as each simulation could only accommodate a single replicate (i.e. one coral with a microsensor at the surface and another microsensor in the bulk seawater).

| **Temp setpoint (°C)** | **Temp setpoint (Δ from summer mean; °C)** | **Realised temp (Δ from setpoint; °C)** | **Standard deviation** |
|---|---|---|---|
| 30 | -2 | -0.005 | 0.128 |
| 30 | -2 | -0.015 | 0.075 |
| 31 | -1 | -0.095 | 0.135 |
| 32 | 0 | -0.025 | 0.155 |
| 33 | 1 | -0.071 | 0.135 |
| 34 | 2 | -0.124 | 0.075 |
| 35 | 3 | -0.048 | 0.189 |
| 36 | 4 | 0.009 | 0.151 |
| 37 | 5 | 0.024 | 0.179 |
| 38 | 6 | -0.035 | 0.218 |
| 39 | 7 | -0.49 | 0.344 |
| 40 | 8 | -1.205 | 0.512 |

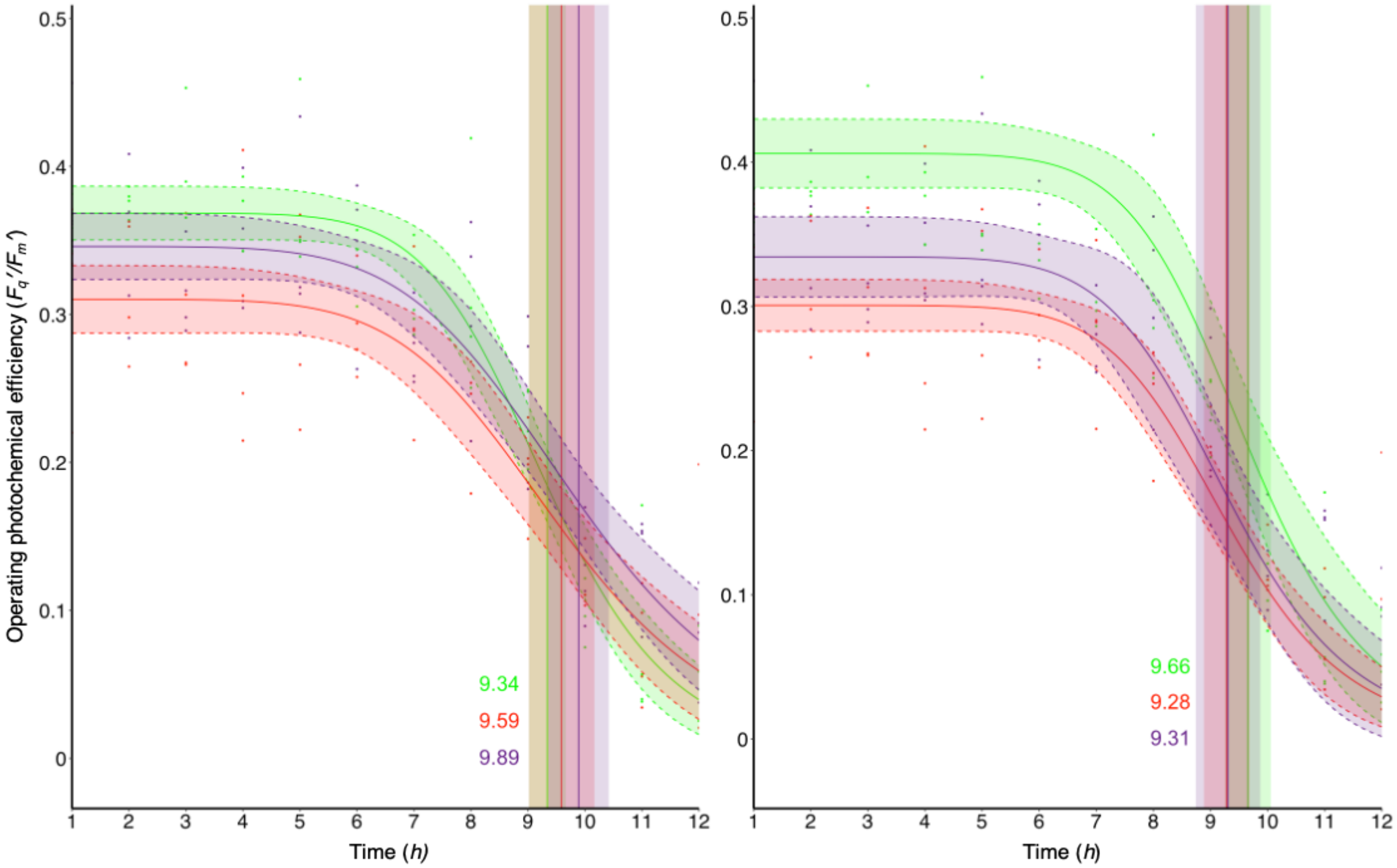


**Supplementary Figure 2**: **Operating photochemical efficiency ($F_q'/F_m'$).** Pulse-amplitude-modulated fluorometry under normal (left) and high (right) flow conditions. All the replicate $F_q'/F_m'$ data for each of the three genotypes used in this study was averaged at each time point and plotted by genotype. Note that each measurement was conducted on each fragment through time, so exposures to time/temperature accumulated throughout the treatment. The values represent the $ET_{50}$ value, at which at 50% drop in photochemical efficiency has been observed. Here, this value represents time (hours) since beginning of simulation of increases in temperature of 1°C per hour, starting at 2°C below the summer mean and ending at 8°C above.

**Supplementary Table 2: Closed-chamber incubations in the absence of heat stress.** 'Time in' represents the start of the experiment. 'Anoxia time' is the time at which dissolved oxygen reached 0 mg/L of $O_2$. 'Time out' is the time after a 4 h exposure to anoxia. After each fragment of *Acropora hemprichii* was removed, it was photographed and observations were recorded before being placed back into a holding tank. After a day in the holding tank, another photo was taken of each fragment. All times are in local time.

| Time in | Anoxia time | Time out | Time out observations | Time out photo | +1 d photo |
|---|---|---|---|---|---|
| 19:30 | 23:50 | 03:50 | Tissue sloughing when moved. Pungent smell when chamber was opened. Unusual patches of pale visible, looks almost like scaring. | 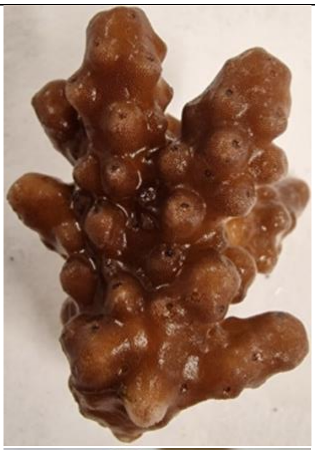 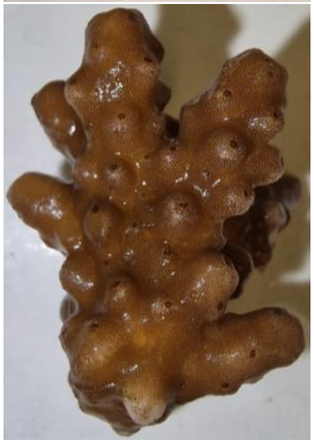 | 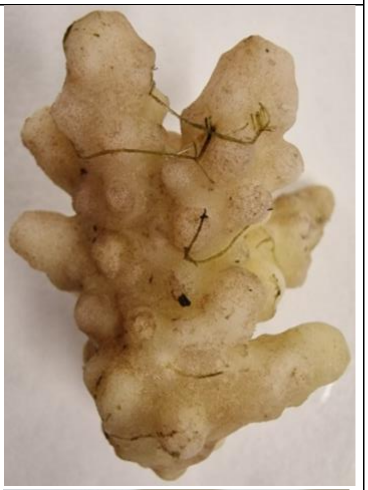 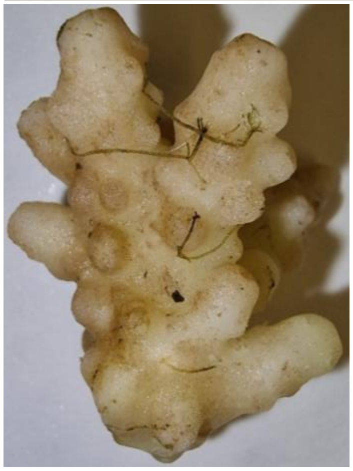 |

| | | | | | |
|---|---|---|---|---|---|
| 19:30 | 23:50 | 03:50 | Tissue sloughing when moved. Pungent smell when chamber was opened. Unusual patches of pale visible, looks almost like scaring. | 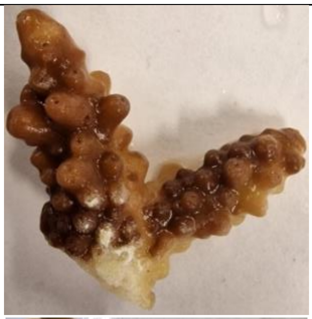 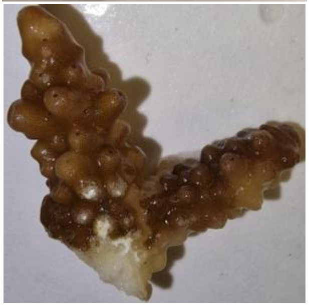 | 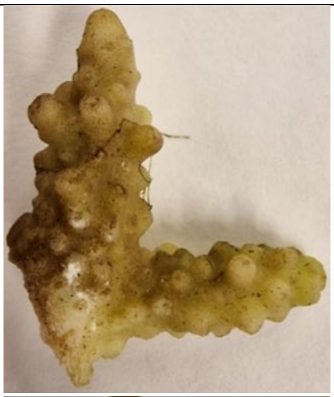 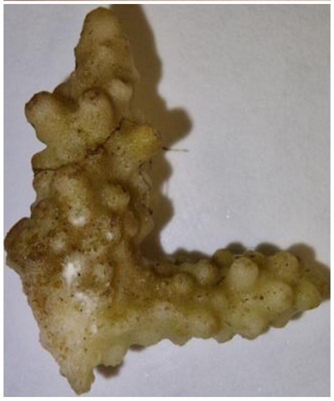 |

| | | | | | |
|---|---|---|---|---|---|
| 19:30 | 01:40 | 05:40 | Tissue sloughing when moved. Pungent smell when chamber was opened. Unusual patches of pale visible, looks almost like scaring. | 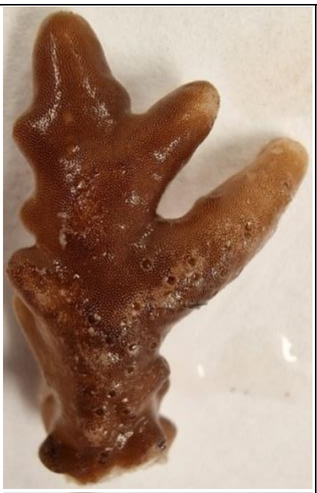 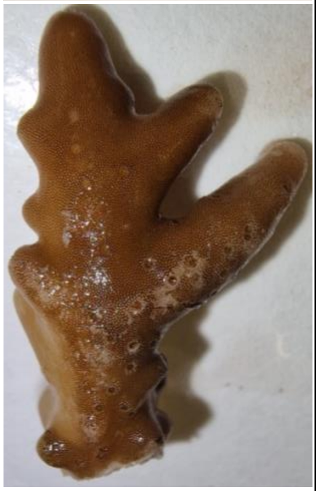 | 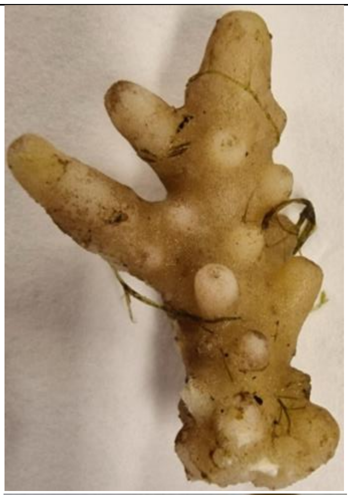 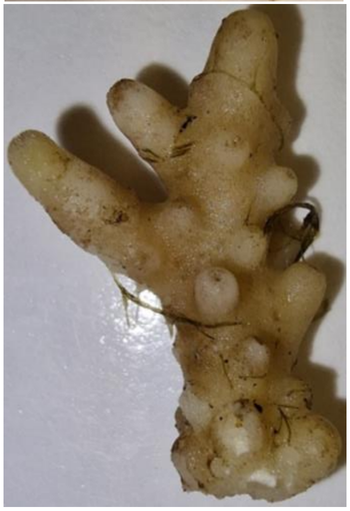 |

| | | | | | |
|---|---|---|---|---|---|
| 19:30 | 01:40 | 05:40 | Tissue sloughing when moved. Pungent smell when chamber was opened. Unusual patches of pale visible, looks almost like scaring. | 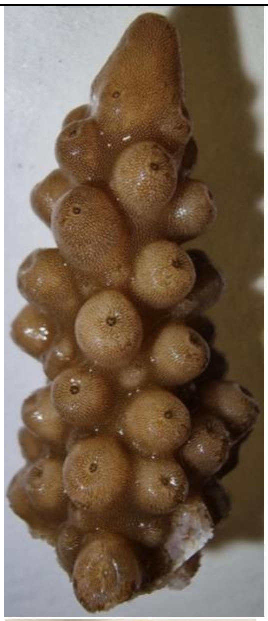 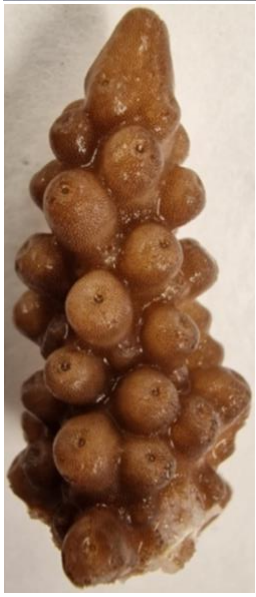 | 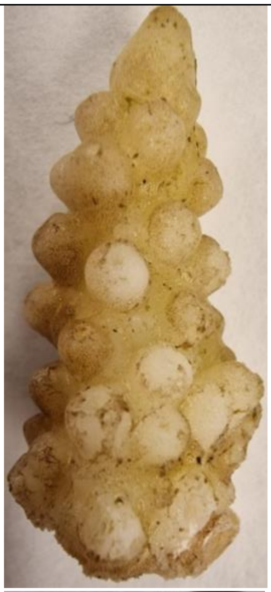 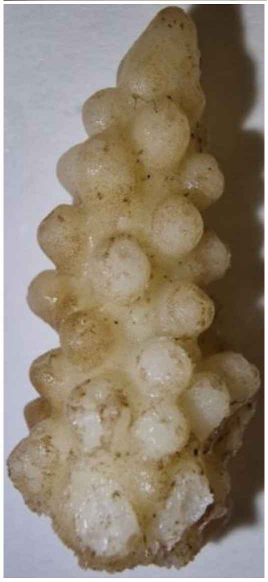 |

| | | | | | |
|---|---|---|---|---|---|
| 19:30 | 01:40 | 05:40 | Tissue sloughing when moved. Pungent smell when chamber was opened. Unusual patches of pale visible, looks almost like scaring. | 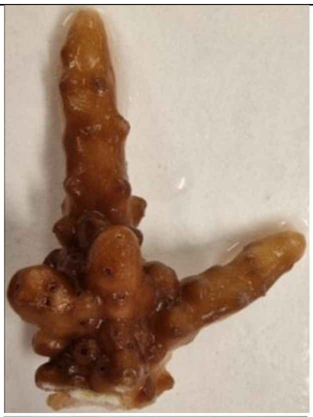 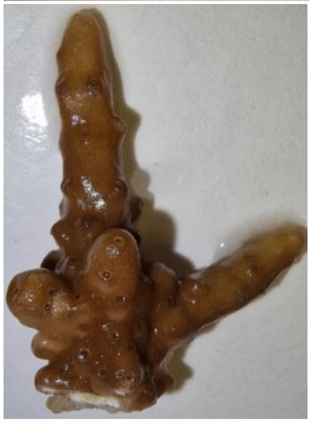 | 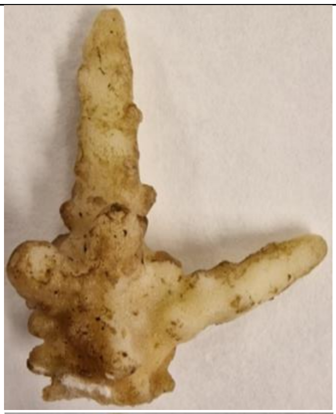 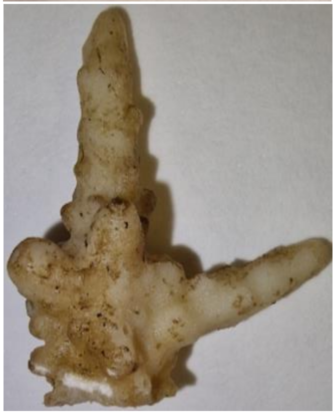 |

| | | | | | |
|---|---|---|---|---|---|
| 19:30 | 02:43 | 06:43 | Tissue sloughing when moved. Pungent smell when chamber was opened. Unusual patches of pale visible, looks almost like scaring. | 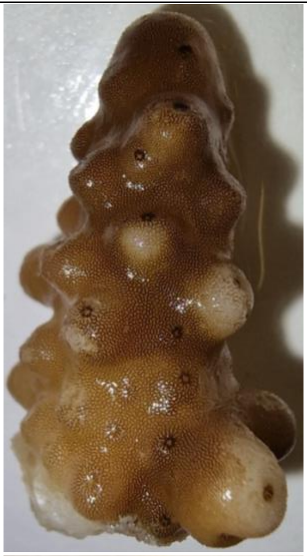 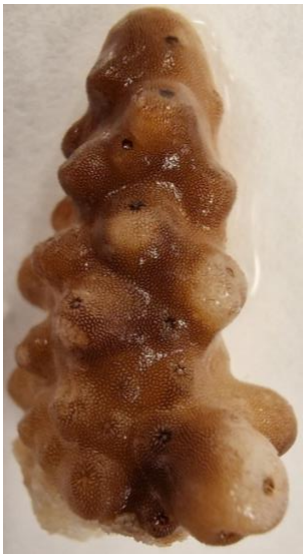 | 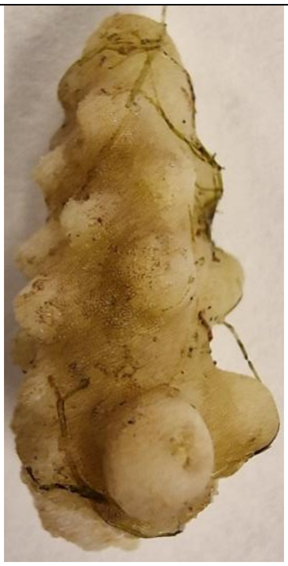 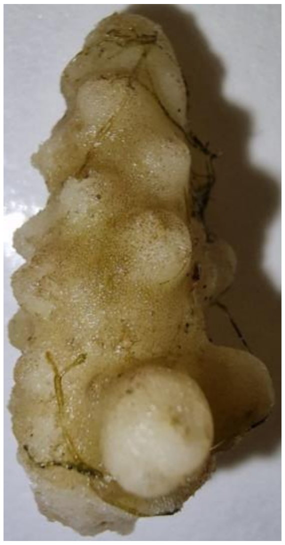 |

| | | | | | |
|---|---|---|---|---|---|
| 19:30 | 02:43 | 06:43 | Tissue sloughing when moved. Pungent smell when chamber was opened. Unusual patches of pale visible, looks almost like scaring. | 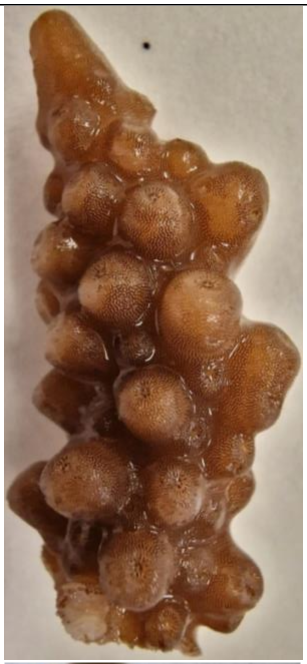<br>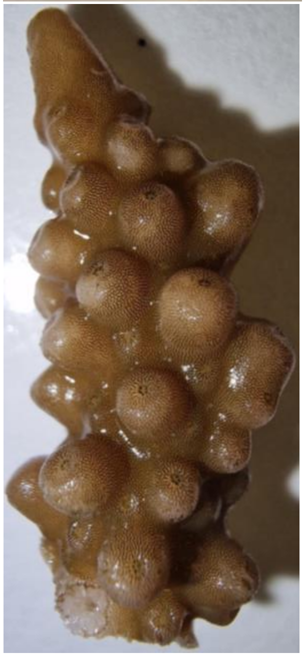 | 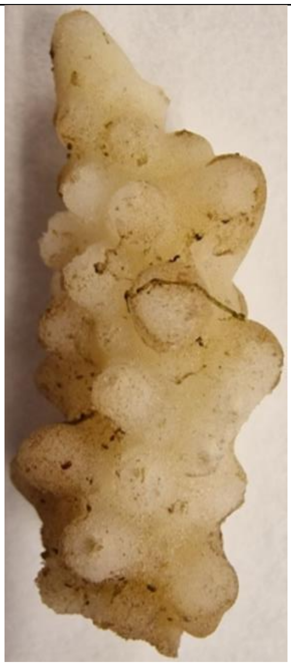<br>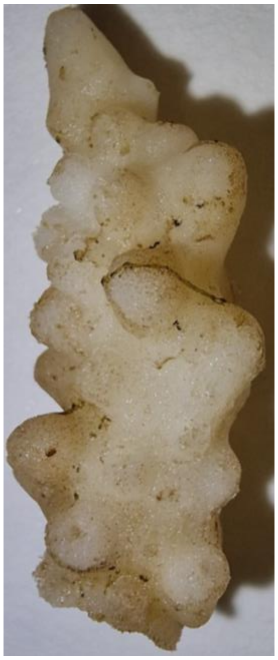 |

| | | | | | |
|---|---|---|---|---|---|
| 19:30 | 02:43 | 06:43 | Tissue sloughing when moved. Pungent smell when chamber was opened. Unusual patches of pale visible, looks almost like scaring. | 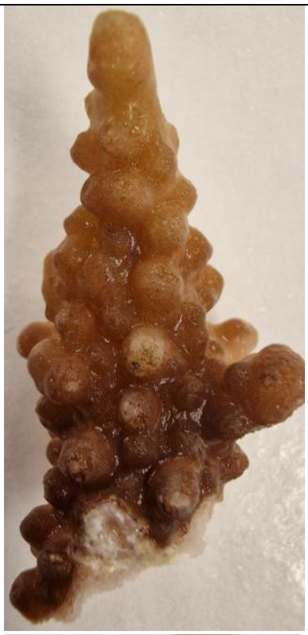<br>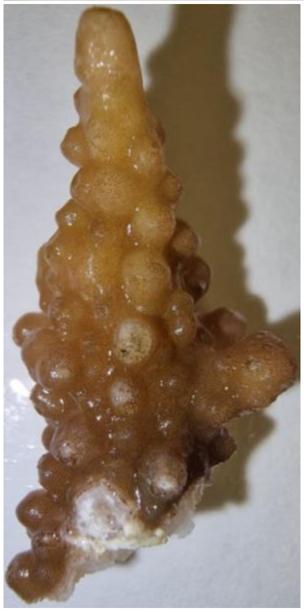 | 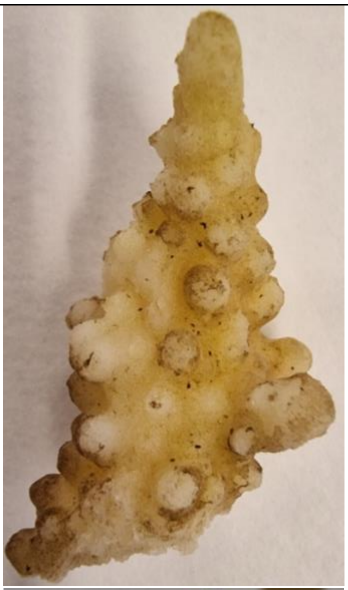<br>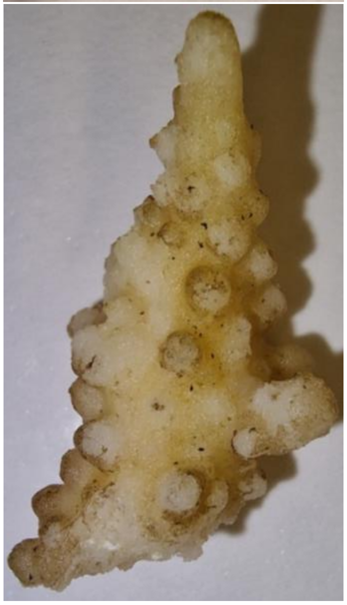 |

| | | | | | |
|---|---|---|---|---|---|
| 19:30 | 04:30 | 08:30 | Small amount of tissue sloughing when moved; nowhere near as severe as the other replicates. Smell not as pungent as other replicates. Surface scarring visible. | 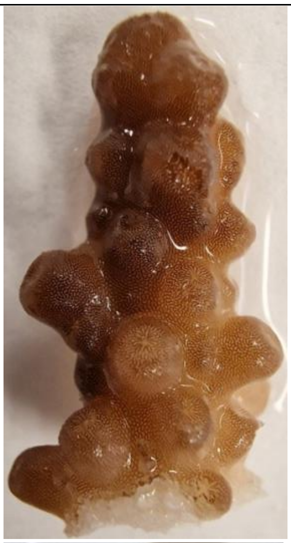 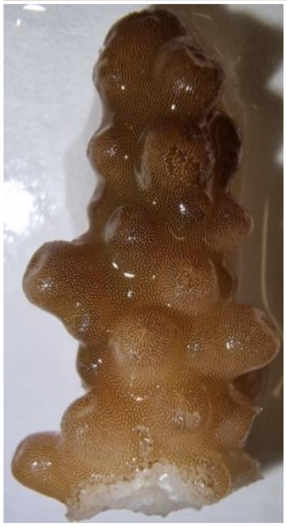 | 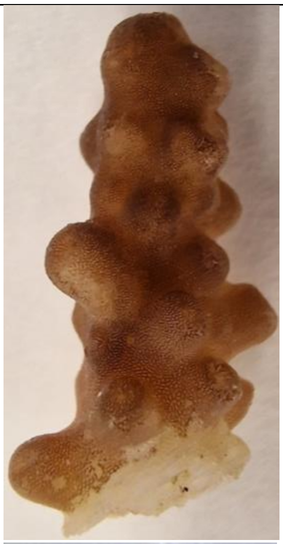 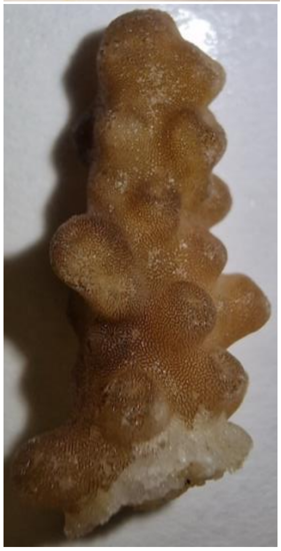 |
